# smiFISH and embryo segmentation for single-cell multi-gene RNA quantification in arthropods

**DOI:** 10.1101/2020.02.29.971390

**Authors:** Llilians Calvo, Matthew Ronshaugen, Tom Pettini

## Abstract

Recently, advances in fluorescent in-situ hybridization techniques and in imaging technology have enabled visualisation and counting of individual RNA molecules in single cells. This has greatly enhanced the resolution in our understanding of transcriptional processes. Here, we adapt a recently published smiFISH protocol (single-molecule inexpensive fluorescent in-situ hybridization) to whole embryos across a range of arthropod model species, and also to non-embryonic tissues. Using multiple fluorophores with distinct spectra and white light laser confocal imaging, we simultaneously detect and separate single RNAs from up to eight different genes in a whole embryo. We also combine smiFISH with cell membrane immunofluorescence, and present an imaging and analysis pipeline for 3D cell segmentation and single-cell RNA counting in whole blastoderm embryos. Finally, using whole embryo single-cell RNA count data, we propose two alternative single-cell variability measures to the commonly used Fano factor, and compare the capacity of these three measures to address different aspects of single-cell expression variability.

## INTRODUCTION

For many years, RNA in-situ hybridization (ISH) and immuno-staining have been the methods of choice for studying gene expression patterns, but have not generally been used to quantify expression levels beyond qualitative differences. This is because signal amplification steps introduce intensity variation and nonlinearity in detection precluding quantitative comparison, and off-target probe or antibody binding can produce substantial false positives^1^. Instead, quantification of gene expression has largely relied on quantitative PCR, microarrays, nanostring technology and bulk RNA-seq. These techniques usually provide only relative expression levels rather than actual RNA numbers, across a pool of cells, so a wealth of information concerning cell to cell variability is lost^2-6^. More recently, single-cell versions of these techniques have been developed, allowing for the first-time quantitation of cell differences in gene expression^7, 8^, known to be critical in influencing single-cell behaviours^9, 10^, differentiation^11^ and disease^12^. However, the spatial context of the cells with respect to both their neighbouring cells, and to the larger tissue or embryo is still lost^13^.

Recently, these limitations have been overcome by the development of single-molecule fluorescent in-situ hybridization (smFISH), which employs multiple short ∼20nt gene-specific DNA probes directly labeled with fluorophores^14, 15^. When multiple short probes bind to target RNA, the single RNA molecules can be visualized and counted as discrete fluorescent spots. Accurate quantification is possible because both false positives and negatives are minimized, since a single off target smFISH probe is below detection limits, and a false negative is unlikely as this would require that most of the ∼40 probes miss the same target molecule. Furthermore, cells remain fixed within the sample rather than being dissociated, so RNA number can be quantified on a cell by cell basis in the spatial and temporal context of the sample. A variant of smFISH was recently developed, in which the gene specific probes have an additional 28nt flap sequence added to the 5’ end, rather than being directly tagged with fluorophore^16^. This flap sequence is identical for all probes in the set. The complementary sequence to the 28nt flap is synthesized with a fluorophore of choice attached to 5’ and 3’ ends, and then prior to use, the complementary flaps are annealed, creating gene specific probes that are now fluorophore-labeled. This simple change in probe preparation vastly decreases cost, since only a single flap sequence is labeled with fluorophore, rather than each unique gene-specific sequence. Accordingly, this approach is termed single-molecule inexpensive FISH (smiFISH).

The original smiFISH publication tests the technique in cultured mammalian cells^16^. In this study, we modify the protocol, and show it to be effective in early and late embryos from five extant and emerging arthropod model species, and also in non-embryonic tissues, specifically *Drosophila* imaginal discs and ovaries. We also test the compatibility of a suite of different commercially available fluorophores, in combination with confocal imaging and a white-light laser, to attain the maximum number of different RNAs that can be visualized simultaneously in the same sample. We combine smiFISH with immunofluorescence for detection of cell membranes, and present a clearly defined analysis pipeline for whole embryo cell segmentation in 3D image stacks, and single-cell RNA quantification for multiple genes. To enable analysis of single-cell variability, we propose a semi-automated method for identifying the immediate neighbours of each cell in the embryo. The Fano factor, (Variance/mean) is commonly used to measure cell variability in expression level^17^, however, due to its limitations, here we offer two alternative measures of variability that better capture individual cell behaviour, and compare the capacity of each method to address different biological questions.

## ONLINE MATERIALS AND METHODS

### Solutions

50% bleach: 50% sodium hypochlorite solution in distilled H_2_O. Embryo wash buffer: 0.1M NaCl, 0.02% triton X-100 in distilled H_2_O. Fix solution: 0.5ml 10X PBS, 0.5ml nuclease-free H_2_O, 4ml ultrapure 10% methanol-free formaldehyde, 5ml heptane. PBT: 1X PBS, 0.05% tween-20, in nuclease-free H_2_O. smiFISH wash buffer: 2X SSC, 10% deionised formamide in nuclease-free H_2_O. smiFISH hybridization buffer: 10% w/v dextran sulphate (molecular weight 6,500-10,000), 2X SSC, 10% deionised formamide in nuclease-free H_2_O. Blocking solution: 1X western blocking reagent (Sigma) in PBT.

### Sample fixation

*Drosophila* embryos collected on apple juice agar plates at 25°C were dechorionated with 50% bleach, alternately washed with distilled water and embryo wash buffer, and shaken in fix solution at 240rpm for 45 minutes. The aqueous solution layer was removed, 10ml 100% methanol added, and shaken for 1 minute to devitellinize embryos. Devitellinized embryos were washed 5x with 100% methanol, then stored in 100% methanol at −20°C. Fixed *Tribolium* embryos were kindly provided by Olivia Tidswell from Michael Akam’s lab. *Tribolium* were dechorionated and fixed as described for *Drosophila*, but for increased devitellinization efficiency, embryos were passed through a 19G needle in ice-cold 100% methanol. Fixed *Nasonia* embryos were kindly provided by Shannon Taylor from Peter Dearden’s lab^35^. *Parhyale* embryos were manually collected from females anaesthetized with 0.01% clove oil in sea water. Embryos were washed 3x with filtered sea water, transferred to fix solution, and shaken at 240rpm for 45 minutes. Embryos were then transferred to a glass dish in 1X PBS, for manual removal of the chorion and vitelline membrane using tungsten needles. Dissected embryos were transferred to a second fixation solution of 4% formaldehyde in 1X PBS for 1 hour, before washing 5x in 100% methanol and storage in 100% methanol at −20°C. For imaginal discs, white pre-pupae were chilled in 1°C 1X PBS, the cuticle opened longitudinally, and pupae fixed in a solution of 4% formaldehyde in 1X PBS for 1 hour (no rocking), before washing 5x in 100% methanol and storage in 100% methanol at −20°C. For ovaries, adult females were thoroughly anaesthetized with CO_2_, placed in 1°C 1X PBT, and ovaries dissected out and opened to expose ovarioles. Dissected ovaries were washed with 1X PBS, then fixed and stored as described above for pupae.

### Probe design

*D.melanogaster* and *D.virilis* mRNA sequences were obtained from Flybase (https://flybase.org); *T.castaneum, N.vitripennis* and *P.hawaiensis* mRNA sequences were obtained from NCBI (https://www.ncbi.nlm.nih.gov/nucleotide/). Complementary 20nt DNA probes against mRNA sequences (up to 48 probes per gene) were designed using the Biosearch Technologies stellaris RNA FISH probe designer tool (free with registration,https://biosearchtech.com). The following sequence was added to the 5’ end of each 20nt probe: CCTCCTAAGTTTCGAGCTGGACTCAGTG. This is the reverse complement of the X flap sequence used in Tsanov *et al*. 2016. Oligos were ordered from Integrated DNA Technologies (IDT), using 25nmole synthesis scale, standard desalting, and at 100uM in nuclease-free H_2_O. The X flap sequence itself CACTGAGTCCAGCTCGAAACTTAGGAGG was 5’ and 3’ end-labeled with CalFluor 540, Quasar 570, CalFluor 590, CalFluor 610, CalFluor 635, Quasar 670, and Quasar 705, and synthesized by Biosearch Technologies. X flap sequence 5’ and 3’ end-labeled with Alexa Fluor 488 was synthesized by IDT.

### smiFISH and immunofluorescence

Fixed samples stored in 100% methanol were transitioned to PBT in stages: 50% PBT, 75% PBT, 100% PBT, 5 minutes per wash. Samples were washed 3x 10 minutes in PBT, then 10 minutes in 50% PBT 50% smiFISH wash buffer, before 2x 30 minute pre-hybridization washes in smiFISH wash buffer at 37°C. Probes were annealed to labeled FLAP sequences according to Tsanov *et al*. 2016. Probe/fluorophore combinations are supplied in Supplementary Table 1. When more than 3 probes were to be used on the same sample, probes were annealed at 5x concentration (20uM), so they could be used at 1/5 normal volume, to avoid large volumes of probe affecting salt and formamide concentration in the subsequent hybridization. Annealed smiFISH probes were diluted in 500ul smiFISH hybridization buffer to a working concentration of 80nM. Probes were hybridized with samples at 37°C for 14 hours. Samples were washed 4x 15 minutes in smiFISH wash buffer at 37°C, then 3x 10 minutes in PBT at room temperature. For immunofluorescence, samples were blocked for 30 minutes in blocking solution, then incubated with anti *Drosophila* alpha-Spectrin (DSHB 3A9) diluted 1:50 in blocking solution for 18 hours at 4°C. Samples were washed 4x 15 minutes with PBT, blocked for 30 minutes, incubated with goat anti mouse Alexa Fluor 488 (ThermoFisher) diluted 1:500 in blocking solution for 4 hours at room temperature, then washed 4x 15 minutes with PBT. In PBT, *Tribolium* and *Parhyale* embryos were manually dissected away from yolk using tungsten needles, imaginal discs were dissected away from pupal carcasses, and single ovarioles and egg chambers were dissected away from one another. All samples were mounted under #1.5 coverslips using prolong diamond antifade mountant with DAPI (ThermoFisher). Due to their size, coverslip spacers were required for *Parhyale* embryos.

### Imaging

Images were acquired on a Leica TCS SP8 AOBS inverted gSTED microscope using a 40x/1.3 or 100x/1.4 HC PL APO (oil) objective. Image stacks for each different species, imaginal discs and ovaries were taken with the following settings: format 2048×2048 or 4096×4096, speed 400Hz unidirectional, sequential line scanning, line averaging 8 or 16, pinhole 1 airy unit. Each channel was gated 1.0-6.0. DAPI excitation 405nm, laser 5%, collection 415-480nm. CalFluor 610 excitation 590nm, laser 20%, collection 600-642nm. Quasar 670 excitation 647nm, laser 20%, collection 657-750nm. *D.melanogaster* embryo image stacks showing all 8 Hox genes, and 5 segmentation genes, were taken with the following settings: objective 40x/1.3, format 4096×4096, speed 400Hz unidirectional, sequential line scanning, line averaging 16, pinhole 1 airy unit. Each channel was gated 1.0-6.0. DAPI excitation 405nm, laser 5%, collection 415-480nm. AlexaFluor 488 excitation 490nm, laser 15%, collection 498-530nm. CalFluor 540 excitation 522nm, laser 15%, collection 530-555nm. Quasar 570 excitation 548nm, laser power 15%, collection 558-575nm. CalFluor 590 excitation 569nm, laser 15%, collection 579-595nm. CalFluor 610 excitation 590nm, laser 15%, collection 605-620nm. CalFluor 635 excitation 618nm, laser 15%, collection 628-650nm. Quasar 670 excitation 647nm, laser 10%, collection 660-680nm. Quasar 705 excitation 670nm, laser 10%, collection 695-780nm. Image stacks were acquired with a 200nm z interval. For images intended for mRNA quantification, z-stack limits were set to just above and below the extent of the smiFISH signal, so that all mRNAs throughout the z-depth were captured. Spectral unmixing of the 8 Hox gene channels was performed in the Leica LAS X v1.8.0.13370 software, using a 30uM radius selection in each channel to build the unmixing matrix.

### Image analysis

Z stacks were stabilized through z to account for any imaging drift, and deconvolved using Huygens Professional v18.04. For single-cell segmentation and mRNA quantification, DAPI, Spectrin, and smiFISH image stacks were combined in Imaris v9.2 software. Spectrin staining forms a clear cell border in z-slices where the cells are in cross-section, but fades out basally at the extent of membrane ingression. The core set of z-slices that do show clear cross-sectional Spectrin staining was identified, and the bottom slice of this core replicated to extend through the full depth of the stack, replacing the basal slices with unclear cell borders. Cells were then segmented in 3D automatically in the Imaris cells module from the Spectrin channel, using a smallest cell diameter of 5um, membrane detail level of 0.5um, and a local contrast filter. Edge cells and any double cells were omitted by filtering the set of detected cells for outliers based on cell volume, sphericity and z-position. smiFISH spots were detected using the spots function, allowing for different spot sizes, with an estimated xy diameter of 0.3um, estimated z diameter of 0.6um, and background subtraction. Spot quality thresholds were set individually for each channel, since brightness and diameter of spots is inherently different between different fluorophores, but for quantitation consistency, these same thresholds must then be used for every embryo analysed. To prepare images for figures, maximum projections of smiFISH channels were generated in FIJI v 2.0.0-rc-49/1.51d. Projections were then combined into RGB images in Adobe Photoshop CS6.

## RESULTS

### Adaptation of smiFISH to arthropod embryos and tissues

smiFISH was originally tested in cultured mammalian cells^16^. Here we applied the smiFISH protocol, with modifications, to embryos of five different arthropod model species – *Drosophila melanogaster* and *Drosophila virilis* (fruit flies), *Nasonia vitripennis* (parasitoid wasp), *Tribolium castaneum* (flour beetle), and *Parhyale hawaiensis* (amphipod crustacean). The evolutionary divergence times of these species is shown in Supplementary Figure 1. We also tested *Drosophila* imaginal discs and ovaries. Our protocol simplifies the original smiFISH buffers, omitting *E. coli* tRNA, BSA and vanadylribonucleoside complex. 1X PBS is swapped for 1X PBT to avoid embryo or tissue clumping, and we also increase the number and duration of washes, to account both for the fact that embryos and tissues are thicker and more complex than cells, and that complete removal of solutions between washes is less feasible.

An identical protocol was used for all species and tissues, the only minor differences were in the sample fixation method, and the final mounting (detailed in online methods). In all species we stained for the same two genes, *even-skipped* (*eve*) in early embryos, and *engrailed* (*en*) in later embryos (Figure 1). Single mRNA resolution was achieved in embryos of all species, with very low non-specific background, evident from the regions outside of stripes that are devoid of signal. In both *Drosophila* species (diverged 50 MYA) *eve* is expressed in seven stripes. Classically, *eve* stripes detected with normal ISH or immuno-staining tend to have a discrete appearance^18-20^, but here magnified panels showing the regions in between stripes at single molecule resolution reveal that *eve* is expressed throughout the entire region enclosed by the seven stripes. The stripes represent waves of alternating high and low expression, with the ‘low’ cells in between stripes containing around 10 mRNA per cell. In accordance with previous observations, *eve* shows different patterns in *Tribolium, Parhyale* and *Nasonia*, which may reflect distinct upstream regulatory inputs, and the differing modes of segmentation in these species compared with *Drosophila*^21-23^.

**Figure 1.**
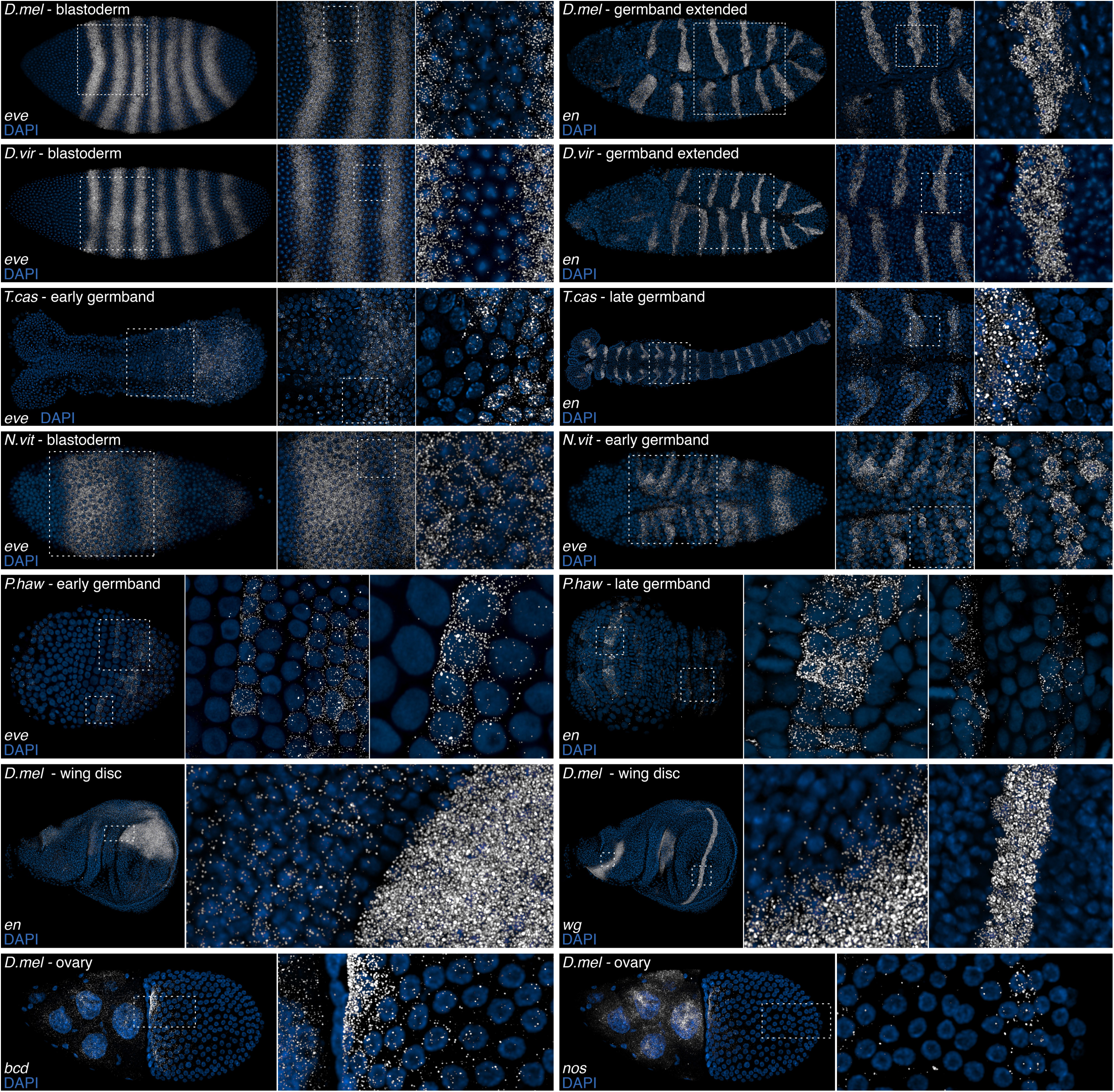
smiFISH in different arthropod species and tissues. smiFISH for the segmentation genes *even-skipped* (*eve*) and *engrailed* (*en*) are shown in early and later embryos from five different arthropod species, *Drosophila melanogaster* (*D.mel*), *Drosophila virilis* (*D.vir*), *Tribolium castaneum* (*T.cas*), *Nasonia vitripennis* (*N.vit*), and *Parhyale hawaiensis* (*P.haw*). Embryos are oriented with anterior to left. smiFISH for *wingless* (*wg*) and *en* is shown in the *D.mel* imaginal wing disc. Ovaries were stained for the maternally loaded RNAs *bicoid* (*bcd*) and *nanos* (*nos*), which accumulate at the anterior and posterior poles of the developing egg respectively. A single egg chamber is shown, oriented with nurse cells and the anterior of the developing egg to left. DAPI was used to stain cell nuclei. All images were acquired using a white light laser scanning confocal microscope with 40X or 100X objectives. White dashed boxes are magnified to the right. Single mRNAs are visible for all samples tested.

Imaginal discs were stained for *en* and *wingless* (*wg*) (Figure 1). Both genes have regions within the wing disc with markedly different expression levels (*en*, magnified panel), and both sharp and diffuse boundaries (*wg*, magnified panels), presumably arising from regional differences in transcriptional regulation. Ovaries were stained for *bicoid* (*bcd*) and *nanos* (*nos*) RNAs (Figure 1). Both genes are highly expressed in the nurse cells. As expected, *bcd* RNAs accumulate at high density at the anterior edge of the developing egg, with a gradient of decreasing concentration towards the posterior^24, 25^. *nos* RNAs are also abundant at the anterior edge, presumably due to proximity to the nurse cells, and as expected, also show some accumulation at the posterior pole of the egg, visible in the magnified panel^26, 27^.

### Simultaneous multi-gene visualization at single molecule resolution

Tsanov *et al*. 2016 show that since smiFISH flaps are first annealed in vitro, probes using the same flap sequence but with different fluorophores can be used together without crossover. Using only the X flap sequence for all smiFISH probe sets, we tested the performance of multiple fluorophores, alone and in combination, with the aim of identifying a maximum set with separable spectra, that would allow simultaneous detection of multiple distinct gene expression patterns at single molecule resolution. We were able to separate nine colours; eight *Drosphila* Hox genes at single molecule resolution together in the same embryo, plus DAPI to stain nuclei (Figure 2). Probe/fluorophore combinations are supplied in Supplementary Table 1. The image is provided as a high resolution supplemental file, where single mRNAs can be observed with zoom. To view eight genes together, optimal excitation and collection from each fluorophore is essential to avoid bleed-through between channels. This image was acquired using an inverted SP8 confocal with white light laser, tunable to each specific excitation wavelength. Narrow collection windows of ∼20nm were set, corresponding to peak emissions of each fluorophore. Line averaging 16x, and high resolution 4096 x 4096 format enabled single RNAs to be resolved. Despite settings that minimized bleed-through, some still persisted between certain channels, so the image was spectrally unmixed following acquisition. To avoid the need for spectral unmixing, a six colour stain using DAPI, AlexaFluor 488, Quasar 570, CalFluor 610, Quasar 670 and Quasar 705 is ideal.

**Figure 2.**
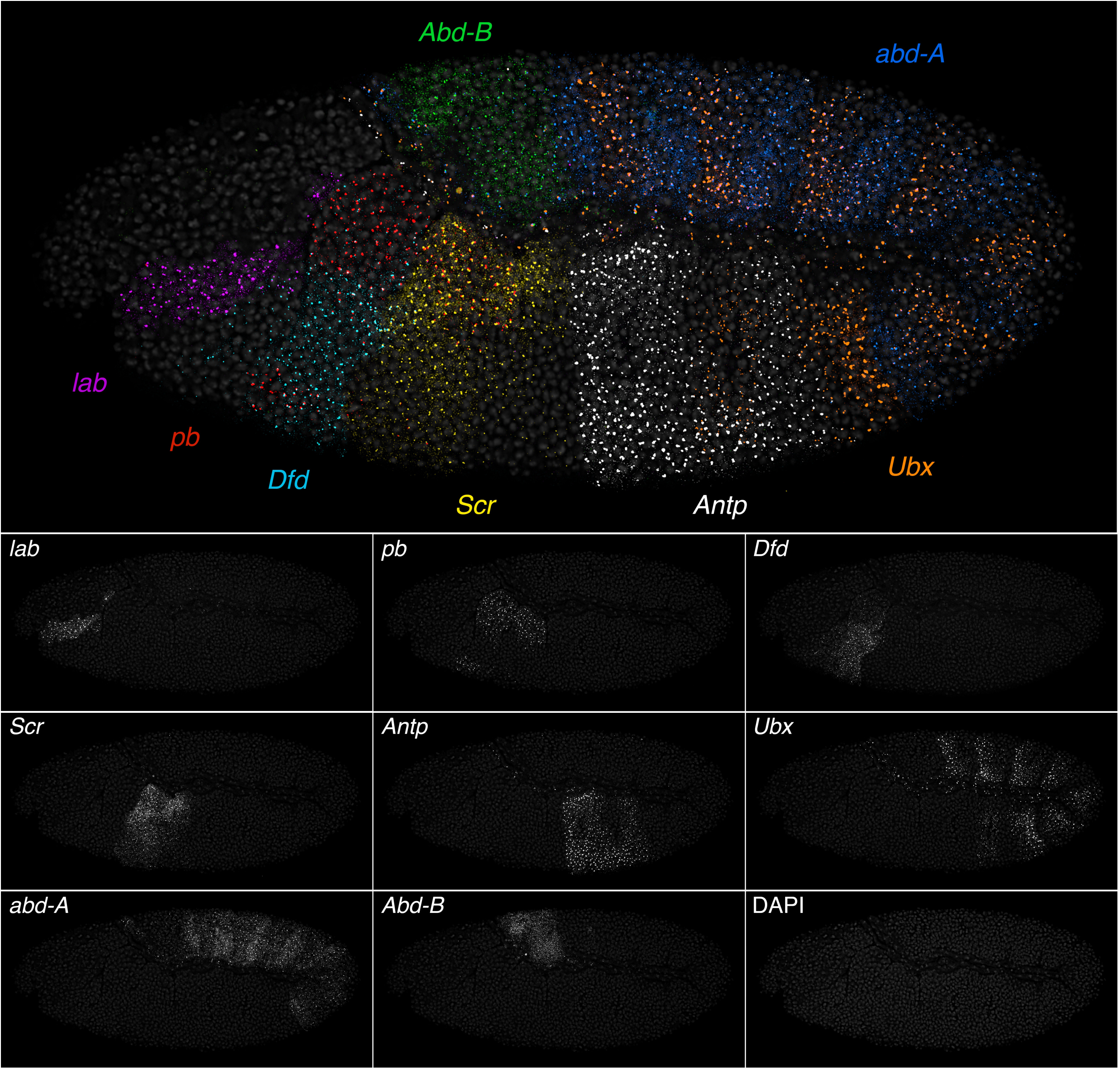
smiFISH and white light laser confocal imaging to visualize all eight *Drosophila* Hox genes at single molecule resolution. A stage 10 germband extended *D.melanogaster* embryo (lateral view, anterior left) with smiFISH staining for all 8 Hox genes, plus DAPI to show nuclei. The X-flap sequence was used for all probes, with the following fluorophores: *labial* CalFluor 610, *proboscipedia* Quasar 570, *Deformed* AlexaFluor 488, *Sex combs reduced* Quasar 670, *Antennapedia* promoter 1 CalFluor 540, *Ultrabithorax* Quasar 705, *abdominal-A* CalFluor 590, *Abdominal-B* CalFluor 635. The image stack was acquired using a Leica SP8 inverted confocal microscope, with 40X objective, and a white light laser, enabling optimal excitation wavelengths for each fluorophore. Peak emissions were captured by narrow ∼20nm tunable collection windows, and the image spectrally unmixed in Leica LAS X software to correct any residual bleed-through. Large bright spots mark accumulations of nascent RNAs at transcriptional sites; smaller fainter spots are single mRNAs. Single mRNAs are most readily visible in the greyscale panels.

### Whole embryo segmentation for single-cell multi-gene RNA quantification

The primary advantage of smFISH is to quantify RNA on a cell-by-cell basis, while preserving positional context. Distinguishing individual cells in culture is straightforward if spacing is sufficiently sparse, but in embryos or tissues is more challenging, and requires a cell membrane marker and segmentation. We tested the compatibility of smiFISH with cell membrane immunofluorescence using a panel of different *Drosophila* antibodies, and found that immunofluorescence is best incorporated after smiFISH, not before. We identified alpha-Spectrin as an ideal marker that clearly defines cell boundaries and is least compromised by the prior smiFISH steps.

To quantify RNAs from multiple genes in single cells, we performed smiFISH in *Drosophila* embryos for four gap genes expressed at blastoderm stage - *hunchback* (*hb*), *giant* (*gt*), *knirps* (*kni*) and *Kruppel* (*Kr*), the pair rule gene *eve*, and marked cell membranes by Spectrin immunofluorescence (Figure 3a). For probe/fluorophore combinations see Supplementary Table 1. Spectrin staining forms a clear cell border in z-slices where the cells are in cross-section (Figure 3b). In cellular blastoderm *Drosophila* embryos, cell membranes are in the process of ingressing between nuclei, but have not yet sealed off the basal side, causing Spectrin staining to fade out basally. mRNAs can be observed at z-planes beyond this basal membrane limit. Therefore, the core set of z-slices that do show clear cross-sectional Spectrin staining was identified, and the bottom slice of this core replicated to extend through the full stack depth, replacing the basal slices with unclear cell borders. The cells module in Imaris software was used to segment the embryo in 3D through the full depth of the z-stack (Figure 3b, middle panel), and the spots function to identify individual RNAs for each gene, which are then automatically assigned to cells (Figure 3b, bottom 2 panels). Details of Imaris analysis steps are provided in online methods. Heatmaps display the number of mRNAs of each gene, in each cell of the embryo (Figure 3c). These illustrate that all five genes show expression domains with graded, rather than sharp borders, consistent with the gap expression patterns being established in response to maternal morphogen gradients such as bicoid, within a syncytial embryo. Separate expression domains of the same gap gene show different overall expression levels, suggesting that transcriptional regulation varies with cell position. Histograms of single-cell data are shown in Figure 3d (cells with zero RNA excluded). For cultured cells, the shape of histogram distributions of this type has been used to make inferences about promoter behaviour^28^. However, this inference methodology assumes that the promoter in each cell has a common behaviour shared throughout the cell population, leading to a certain signature evident from the distribution. For example, a promoter with high bursts of transcription followed by long off periods is expected to produce a distribution with a long tail to high values^28^, similar to that found here for *eve*. However, it is important to note that such inferences cannot be made for non-ubiquitously expressed genes in whole embryos and tissues, as the assumption does not hold true. The distributions in Figure 3d represent a mixed population of cells, where each gene shows a variety of different transcriptional behaviours, depending on spatial position within the embryo. The resulting variety in histogram shapes is therefore just a reflection of the different gene expression patterns, not the product of a common promoter behaviour. To illustrate this point, compare the histograms for *Kr* and *eve. Kr* is primarily expressed in a single broad stripe. The low proportion of cells with 15-65 RNA represents the swift spatial transition between very low expressing edge cells, and high expressing cells within the stripe, while the bump between 65-130 corresponds to the large number of high expressing cells within the stripe. In contrast, *eve* is expressed in seven narrow stripes. Multiple stripes means more edges, so a high proportion of cells have intermediate RNA numbers, filling out the 6-65 bins, and a lower proportion of high expressing cells in the centre of stripes, so loss of the bump between 65-130. The histogram shapes of these genes are therefore explained by their patterns, and are not an emergent property of a consistent promoter behaviour.

**Figure 3.**
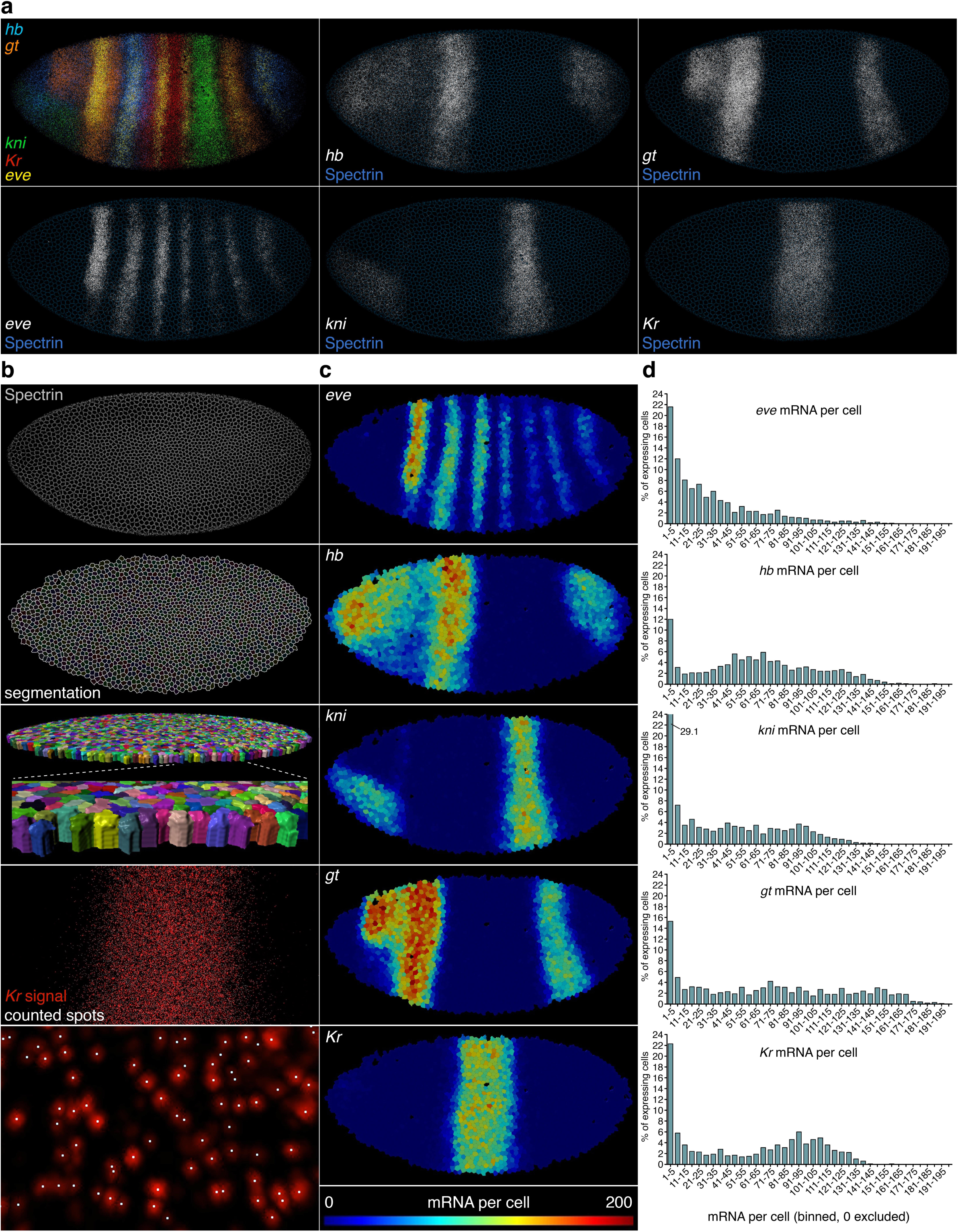
smiFISH with membrane immunofluorescence allows whole embryo 3D segmentation and multi-gene single-cell RNA quantification. **a**) Stage 5 cellular blastoderm *D.melanogaster* embryo (lateral view, anterior left) with maximum projections of smiFISH for the pair rule gene *even-skipped*, and four gap genes: *hunchback, knirps, giant* and *Kruppel*. Nuclei are stained with DAPI, and cell membranes stained by immunofluorescence, using mouse anti *Drosophila* Spectrin, and goat anti mouse AlexaFluor 488. **b**) The cells module in Imaris 9.2 software was used to automatically segment Spectrin staining in 3D through the confocal stack, creating individual cell volumes. The Imaris spots module was used to automatically identify mRNA spots for each gene; and automatically assign spots to cell volumes based on x,y,z coordinates. **c**) Heatmaps displaying mRNA number per cell for each of the five genes. **d**) Histograms of mRNA number per cell for each gene, using bins of five with zero excluded. The shape of histogram distributions is a product of the expression patterns of the genes.

### Semi-automated cell neighbour detection for cell to cell variability analysis

To assess variability in gene expression, one can analyze the mean and spread of RNA values within the whole population of cells, and compare individual cells to this distribution. However, since a given gene may show complex patterns comprising different domains expressing at different levels, analyzing cells together as a single pool may not be informative. Single-cell variability is better addressed by comparing variability between a cell and its immediate neighbours. We define immediately neighbouring cells as those that directly share a membrane border in the Spectrin channel. Using a 2D segmentation plane from the embryo shown in Figure 3b, immediate neighbour number of each cell was manually counted in half of the embryo (Figure 4a). A range of 2-8 immediate neighbours per cell was found, and the frequency distribution is shown in Figure 4b. A neighbour-finding python script was written, using the function scipy.spatial.distance_matrix(coords, coords) from the package scipy.spatial, to calculate the distance between the centre point coordinates of every cell, using the Eucledian distance: √ (x_1_-x_n_)^2^ + (y_1_-y_n_)^2^. The distance matrix was then filtered setting an upper threshold corresponding to a specified radius, such that only cells within that radius are considered neighbours (Figure 4c). To find the correct radius value, we ran the neighbour finding algorithm using different radii, and compared the resulting neighbour number distributions to the manual count (Figure 4d vs 4b). A radius of 8.2 returned a histogram almost identical to the manual count, confirming this as an optimal radius to identify just the immediate neighbours of each cell.

**Figure 4.**
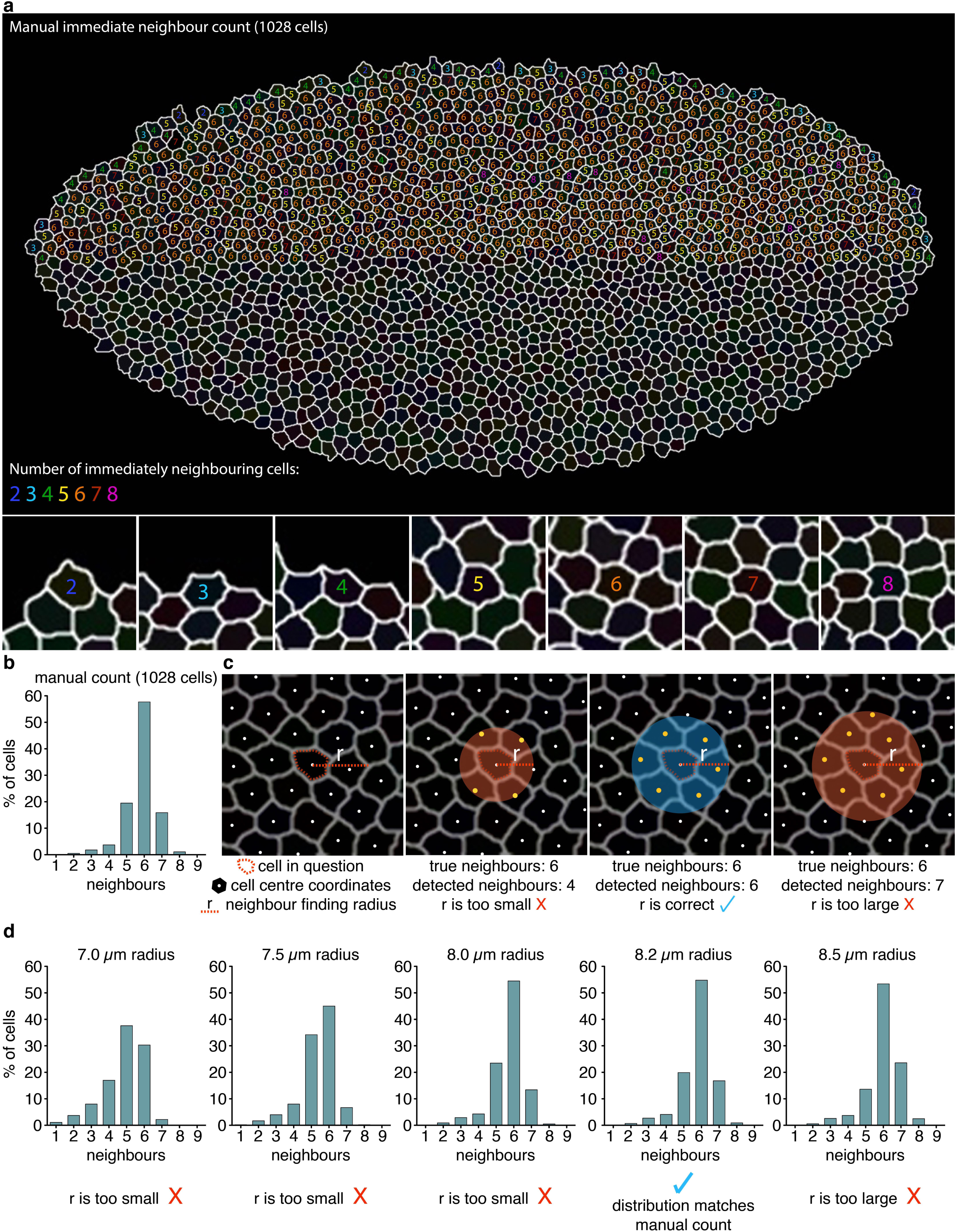
Semi-automated identification of immediately neighbouring cells for single-cell variability analysis. **a)** 2D segmentation plane from a cellular blastoderm *D.melanogaster* embryo with anti-Spectrin membrane staining. The number of immediately neighbouring cells (defined as directly sharing a portion of membrane) was manually counted, for each cell in half of the embryo. A range of 2-8 immediate neighbours was found. **b**) Histogram summarising the manual count, showing the percentage of cells that had each number of immediate neighbours. **c**) Depiction of the neighbour finding concept. The centre XY coordinates of each cell were compared to every other cell in the embryo to calculate the distance between the centres of each cell. A neighbour finding radius is set, such that only cells within that given radius are returned. A correct radius value is essential to accurately identify only and all immediate neighbours. **d**) The neighbour finding script was run using different radii, and the resulting histograms compared to the manual count histogram. 8.2um was identified as optimal, returning a histogram almost identical to the manual count (b).

### New measures to capture numerical and proportional single-cell variability

The smiFISH panels in Figure 5b show *eve*-expressing cells from an early germband *Parhyale* embryo, and highlight how a single cell can have a markedly different expression level from its immediately adjoining neighbours. Such single-cell variability within a population has been shown to have important biological relevance, for example in fate determination^11^, cell behaviour^9^, and disease^12^. Fano factor is a commonly used measure of local mRNA variability, and is calculated as Variance/mean (Figure 5a). Variance and mean are population measures, so all cells in the group are assigned the same Fano value, the resolution of which is therefore dictated by the radius of the neighbour finding. However, Fano factor cannot distinguish single variable cells *within* the neighbour group. This is illustrated by comparing the three hypothetical scenarios depicted in Figure 5 c, d & e. The centre cells in panels c and e are equally different from their neighbours, both proportionately and numerically, whereas the centre cell in panel d is not very variable, being the same as all but one of its neighbours. However, Fano factor fails to distinguish any difference between c and d (both 6.11) and incorrectly finds e much more variable than c (42.37 vs 6.11). To overcome this limitation, we devised two alternative variability measures, the local numerical cell variability (NV), and the local proportional cell variability (PV) (Figure 5a). Both measures express how different an individual cell is from its immediate neighbours. NV is normalized by the maximum mRNA per cell for the whole cell population, therefore a high NV value highlights cells whose mRNA difference from their immediate neighbours is numerically large in terms of the maximum level at which that gene can be expressed. PV is normalized by the maximum just for the neighbour group, and so high PV does not necessarily mean a large difference in actual mRNA number, just that the cell has a high proportional difference from its neighbours. Both measures return values between 0 (no variability) and 1 (maximum variability). In the scenarios shown in Figure 5c-f, 550 is used as the population maximum. Both NV and PV find the centre cells in scenarios c and e to be equally variable, and d to be less variable. Importantly, NV is the same between scenarios c, e and f, since the numerical RNA difference between the centre cell and each neighbour is the same, whereas PV finds scenarios c and e to be proportionally more variable than f.

**Figure 5.**
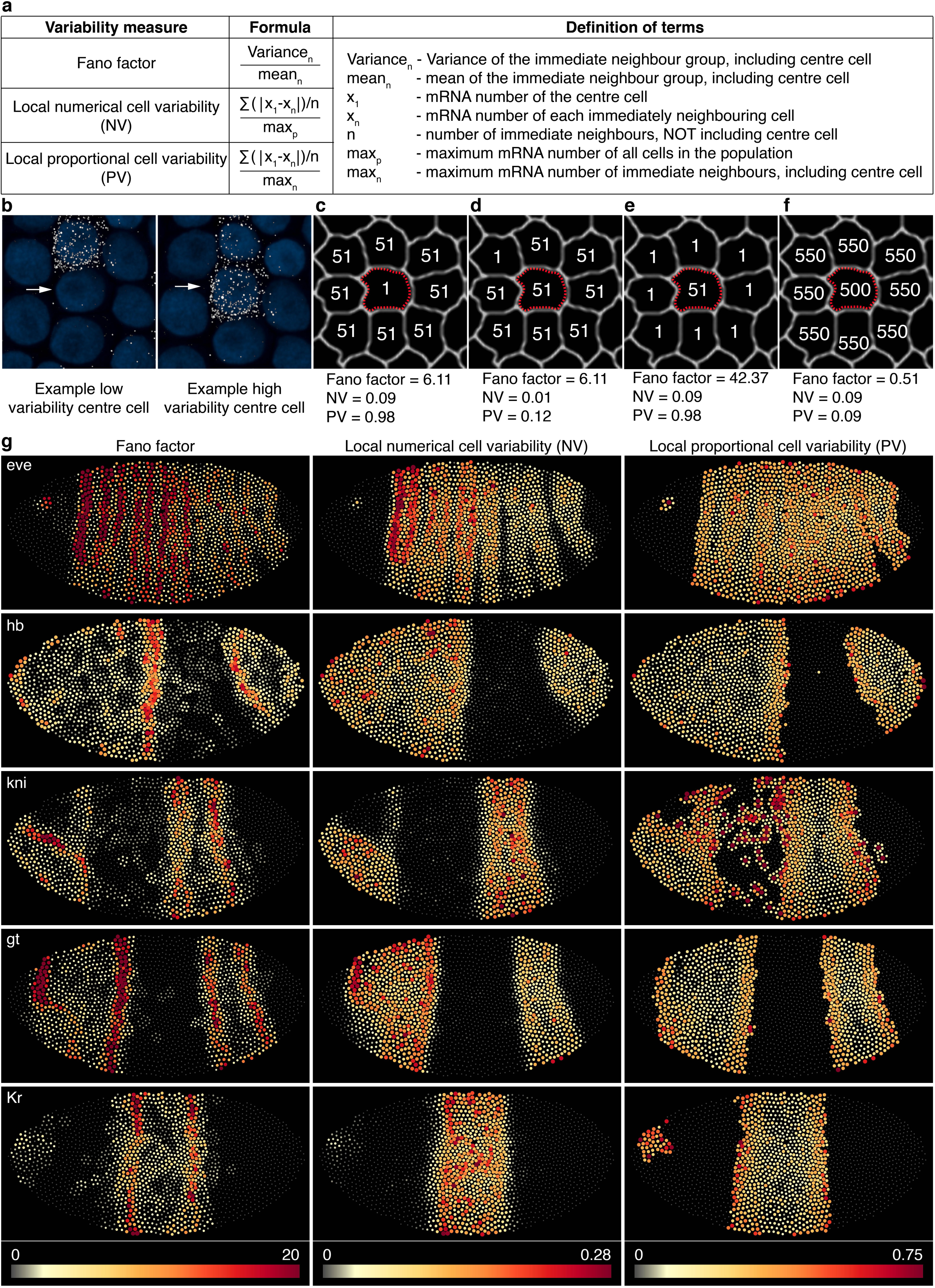
Three alternative measures to express cell to cell mRNA variability. **a**) Formulae for Fano factor, a commonly used measure of cell variability, and two alternative measures designed to better capture individual cell variability. **b**) Cells from the anterior stripe of *eve* expression in *P.hawaiensis* early germband embryo, highlighting that individual cells can differ greatly in expression from their immediate neighbours. The centre cell (white arrow) in the left panel is similar to all its neighbouring cells except one, whereas the centre cell in the right panel is highly different from all of its neighbours except one. **c-f**) Hypothetical scenarios of neighbour group mRNA variability, to highlight the capacity of each formula to capture the variability of the single centre cell. 550 mRNA/cell is used as the population maximum. Fano factor incorrectly returns the same value for c and d, and incorrectly finds e to be more variable than c. NV correctly returns the same value for c, e and f, and a low value for d. PV correctly returns the same value for c and e, and lower values for d and f. **g**) Heatmaps show the three different variability measures, calculated for each cell in the embryo for five segmentation genes, *even-skipped, hunchback, knirps, giant* and *Kruppel*. Dots representing cells are scaled in both size and colour by the variability value.

Fano factor, NV and PV were calculated for the five genes shown in Figure 3, using the optimum immediate neighbour finding radius of 8.2um (determined in Figure 4). Variability scores are displayed as heatmaps (Figure 5g). Cells outside of expression domains that have a single mRNA, surrounded only by non-expressing neighbours, have the maximum PV score of 1. While this is correct, we were more interested to highlight cells that had high PV within actual expression domains. Therefore when calculating PV, cells were filtered on the criteria of neighbour group mean >1; cells failing this criterion were assigned a score of 0. The heatmaps show how Fano, NV and PV highlight different aspects of variability. Fano picks out the edges of expression domains. It acts like a moving average variability, and therefore highlights the regions (but not individual cells) where RNA number is changing the most with position. Within the centre of expression domains, Fano is generally low, suggesting a similar expression level. In contrast, NV can highlight individual cells within the centre of domains that have a high difference in mRNA number from neighbours; cells that were overlooked by the Fano factor. For example, contrast NV and Fano for *kni* and *Kr*. PV highlights cells that are proportionately most different from neighbours, which tends to be cells at the extreme edges of domains, at the transition between off and on. However, individual cells with high PV can still be observed throughout expression domains of each gene.

## DISCUSSION

Whole genome DNA and RNA sequencing is becoming increasingly feasible and affordable, and consequently the number of non-model organisms with whole or partial genome sequence is rapidly growing. Since only ∼1kb of gene sequence is required to design a probe set, smFISH can be applied with ease to non-model species, revealing both expression patterns and levels. Here we have tested smiFISH, with modifications, across a range of arthropod species and sample types, and found that it enabled single mRNA visualization with consistency and high specificity. We also combined smiFISH with subsequent membrane immunofluorescence, allowing whole embryo single-cell segmentation. The anti *Drosophila* alpha-Spectrin antibody used did not work in the non-*Drosophilid* species tested, so appropriate species-specific membrane antibodies are required for use in different organisms.

smiFISH makes multiplexing simple and flexible, and therefore imaging becomes the limitation on how many genes can be viewed together. Using an imaging strategy to optimize fluorophore excitation and capture of emission peaks, we could image nine different channels simultaneously (with spectral unmixing), or six channels without unmixing. The capacity to image more genes simultaneously is advantageous as it allows more potentially interacting genes to be studied within the same cells, thus eliminating error due to sample variability.

A major strength of smFISH is that position of the cell within the sample is preserved, which allows variability to be analysed on a cell by cell basis. We compared a commonly used measure of cell variability, the Fano factor, with two alternative measures termed NV and PV, that were devised to better highlight individual cell variability. Each measure has its own strengths and limitations, and therefore is appropriate for different applications. The Fano factor highlights regions where the RNA number is changing most with cell position, but was not capable of comparing individual to their immediate neighbours. In contrast, NV was effective at highlighting individual cells that were markedly different numerically in mRNA from their neighbours. NV is therefore a relevant measure for questions where the absolute RNA number is important, for example when a threshold expression level is required for a particular process to occur^29, 30^, or post-transcriptional buffering mechanisms that maintain constant mRNA levels^31, 32^. PV also effectively highlighted individual cells that differed from their neighbours, but in proportional expression rather than actual. The PV measure is most relevant for questions involving the mechanisms of RNA production, such as promoter dynamics, and the effects of enhancers and transcription factors. Large PV values may indicate fundamentally different transcription dynamics between cells. This is not necessarily true of high NV, which could be attained in a region of high expressing cells all displaying the same fundamental promoter behaviour, but with some stochasticity that causes a proportionally small, but numerically large RNA difference between cells^33, 34^.

In summary, this work provides a straightforward methodology applicable across a variety of different animal systems, enabling in-depth molecular analyses that traditionally were only feasible in established model systems. Our analysis pipeline to obtain single-cell RNA counts in whole embryos is relevant for studying diverse aspects of expression analysis, and we anticipate that the detailed multi-colour imaging strategy provided here will prove valuable for analysis of gene networks. Finally, it is our view that methods to appropriately analyze spatial cell to cell variability will yield a new level of information critical to understanding how individual cell behaviors lead to biological outcomes.

## ACKNOWLEDGEMENTS

Special thanks to Olivia Tidswell from Michael Akam’s lab, and to Shannon Taylor from Peter Dearden’s lab, for kindly providing us with *Tribolium* and *Nasonia* embryos respectively, and to Peter March from the Manchester bioimaging facility. Thanks to Hilary Ashe and Sam Griffiths-Jones for valuable comments. Attendance to the summer embryology course at Marine Biological Lab, Woods Hole, encouraged the conception of this work, so we are grateful to Rich Schneider and Dave Sherwood for their work in running the course.

## AUTHOR CONTRIBUTIONS

Conceptualization L.C, M.R and T.P, experiments L.C and T.P, imaging and image analysis T.P, analysis formulae L.C and T.P, coding and data analysis L.C, writing – original draft L.C and T.P, writing – review & editing L.C, M.R and T.P, funding acquisition M.R and T.P.

---

**Supplementary Table 1.**
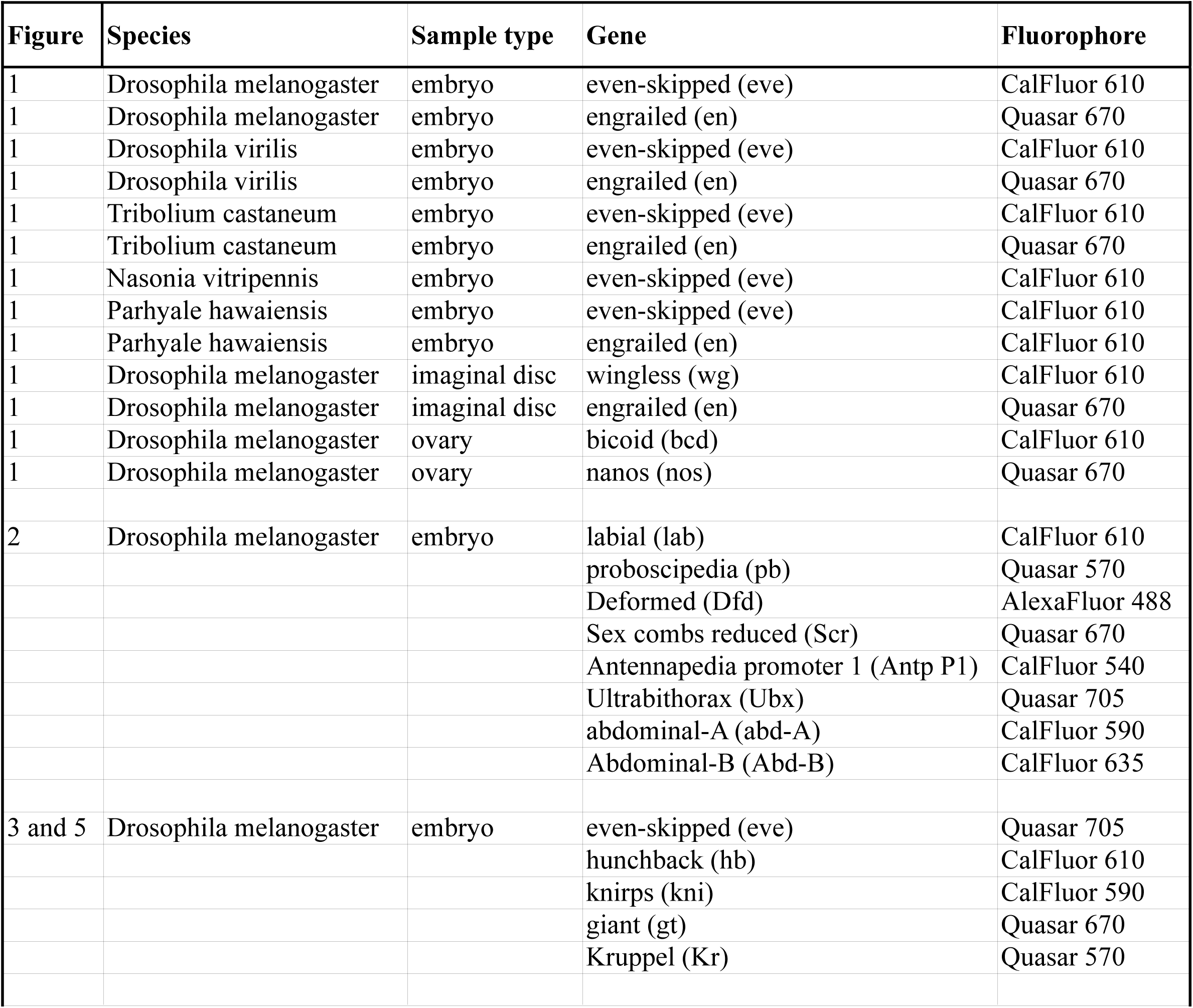
All probe-fluorophore combinations used in this study.

**Supplementary Figure 1.**
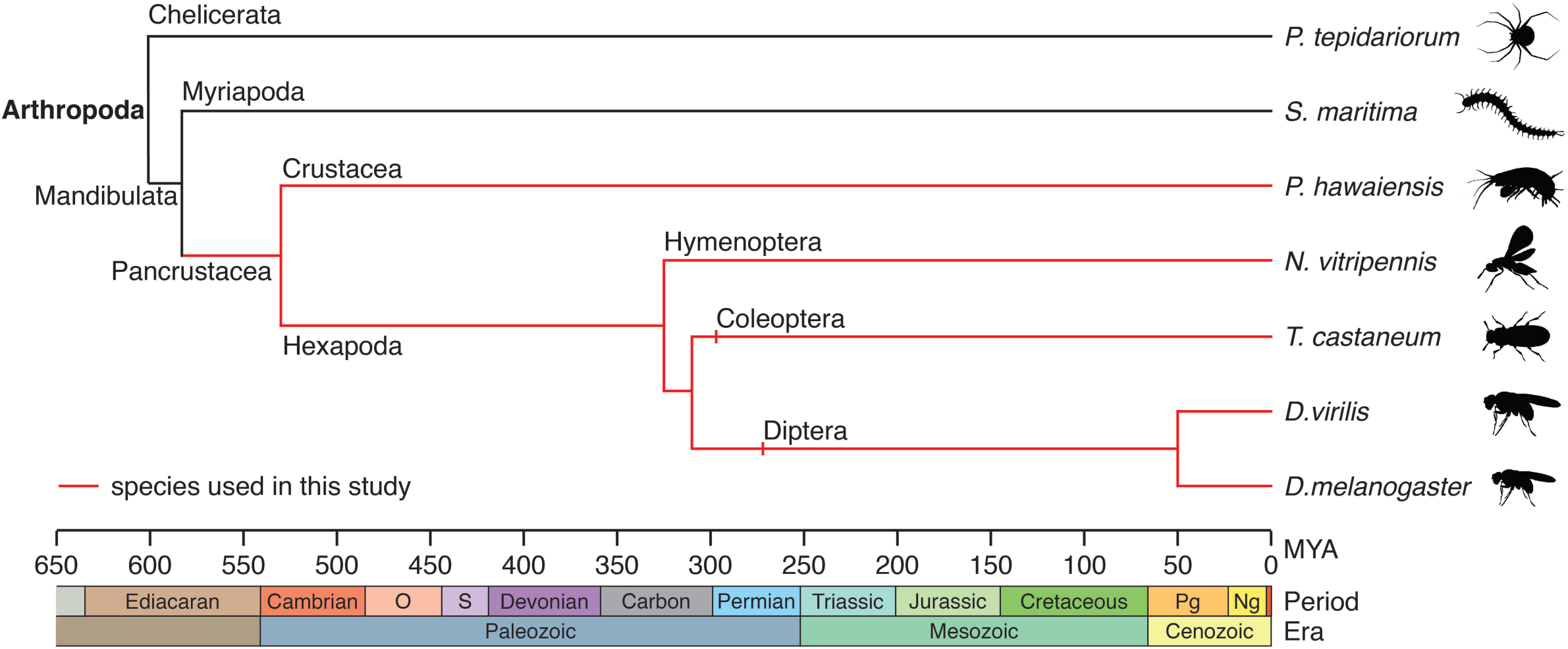
Evolutionary divergence times of different arthropod model species. The species used in this study are highlighted in red, and belong to the clade pancrustacea, which emerged ∼530 MYA, and comprises all hexapods and crustaceans.

